# *In vivo* deep-brain microscopy at submicrometer resolution with refractive index-matched prism interfaces

**DOI:** 10.1101/2025.05.30.657010

**Authors:** Taiga Takahashi, Yuanyuan Zhou, Motosuke Tsutsumi, Chihiro Ito, Azumi Hatakeyama, Hirokazu Ishii, Akiyoshi Saitoh, Hiroshi Yukawa, Junichi Nabekura, Tomomi Nemoto, Kohei Otomo, Naoji Matsuhisa, Masakazu Agetsuma

## Abstract

The mammalian brain is a thick and densely layered structure comprising a huge number of neurons that work together to process information and regulate brain functions. Although various optical methods have been developed to investigate deep brain dynamics, they are limited by technical constraints, invasiveness, suboptimal spatial resolution, and/or a restricted field of view. To overcome these limitations, we developed an implantable, optically optimized microprism interface with a refractive index matched to that of brain tissue and water, enabling minimally-invasive, wide-field two-photon imaging method with enhanced brightness and sub-micron resolution in deep prefrontal areas.

## Main

Neurons throughout the brain communicate with each other via submicron-scale structures called synapses, jointly processing information and contributing to cognition, learning, and behavior^1-5^. *In vivo* two-photon imaging of the mouse brain has yielded unprecedented insight into neural network functions due to its high spatial resolution. This method can be used to record neural activity from wide brain areas at cellular resolution^6-9^, uncovering the mechanisms underlying the computations performed by hundreds to thousands of neurons^4, 5, 10-13^. Two-photon imaging also enables longitudinal monitoring of neural structures during brain development, motor skill improvement, perceptual modifications, and learning^14-21^. However, two-photon imaging has an approximately 1.0-mm depth limitation^22^. In mice, two-photon imaging of deeper brain areas, such as the medial prefrontal cortex (mPFC), located approximately 2 mm or deeper beneath the dorsal surface of the brain and crucial for higher-order cognitive functions^4, 23^, is technically challenging.

Optical imaging techniques using a gradient index (GRIN) lens provide access to deeper brain regions proportional to the length of the lens and can achieve a field of view (FoV) ranging from 0.2 to 0.8 mm^2^, depending on the lens diameter^24, 25^. This approach, however, requires creating a lesion above the imaging site, and deeper access necessitates a larger lesion—an undesirable trade-off given the integrative nature of brain function. This is especially true in the case of the mPFC, where distinct subdivisions interact in cooperative or competitive roles to regulate associative fear memory processing^26^. The depth of view obtained through the GRIN lens is also limited by its optical characteristics. To avoid brain lesions, a three-photon imaging technique was developed and is effective for optically accessing depths of up to 1.1 mm, enabling long-term imaging of neuronal structures and activity^27, 28^. However, spatial resolution in deep regions remains limited due to optical aberrations and tissue scattering. Although Recent advances incorporating adaptive optics (AO) have demonstrated improved spatial resolution beyond 1.4 mm depth^29^, these setups require advanced hardware and complex calibration procedures. Consequently, despite their advantages, these three-photon-based imaging methods are not yet as widely accessible as two-photon microscopy for *in vivo* deep-brain imaging.

Recently, a technique based on a right-angle glass microprism was developed for minimally invasive optical access to deep brain areas^4, 23, 30-33^. By inserting a glass prism along the midline or other deep fissures and placing it tightly beside the target brain region, nerve transections can be avoided. Two-photon imaging through the microprism enables neural activity recordings from hundreds of neurons in mouse mPFC with a wide FoV (2×2 mm^2^), providing insight into how neural populations encode associative fear memory^4^. The effectiveness of microprism-based deep-brain imaging has also been demonstrated in other brain areas associated with higher brain functions, including the posterior mPFC^23^, medial entorhinal cortex^30, 31^, and insular cortex^32, 33^.

One remaining imaging challenge when using a conventional glass microprism is the refractive index mismatch between the brain tissue (1.35–1.40) and the glass (1.52) (Fig.1a)^34^. This mismatch, caused by the presence of the glass prism within the optical path of the objective lens, induces spherical aberrations and reduces the spatial resolution and number of collected photons. Lower brightness also decreases the signal-to-noise (S/N) ratio, often requiring longer exposures or more averaging, which consequently reduces temporal resolution.

In the present study, we developed an optically advanced prism interface, named the Prism with Refractive Index Matched Interface to SpeciMen (PRIMISM), which is designed to match the refractive index of brain tissue, reduce spherical aberrations, and enable bright and high-resolution observations in deep brain tissue. We confirmed that the PRIMISM significantly improved spatial resolution and brightness in microprism-based deep-brain imaging. Using the PRIMISM, we established a platform for high-resolution structural and functional *in vivo* imaging across nearly the entire mouse mPFC, suggesting its potential applicability to other deep brain regions.

## PRIMISM improves spatial resolution and brightness

The PRIMISM was specifically designed to minimize spherical aberrations and enhance fluorescence photon collection efficiency, resulting in significantly improved image brightness and spatial resolution compared to conventional glass microprisms. The PRIMISM (each face 2×2 mm^2^, the same as conventional glass prisms) was engineered with a refractive index that closely matches that of water (1.33) and brain tissue (1.35-1.40)^35-37^, which is crucial for minimizing the effects of aberrations and enhancing image quality in deep brain regions (Fig. 1a,b). The interface of the PRIMISM is nearly invisible when it is embedded in an agarose gel because the PRIMISM has almost the same refractive index value as water (Fig. 1c). Detailed fabrication procedures are provided in Supplementary Fig. 1, and dimensional specifications are shown in Supplementary Fig. 2. The fabrication process employed a technique to eliminate the impact of the bubble, which is unavoidably generated within the PRIMISM during UV curing, from the optical path (for excitation and emission) by adjusting the mirror position.

**Figure 1.**
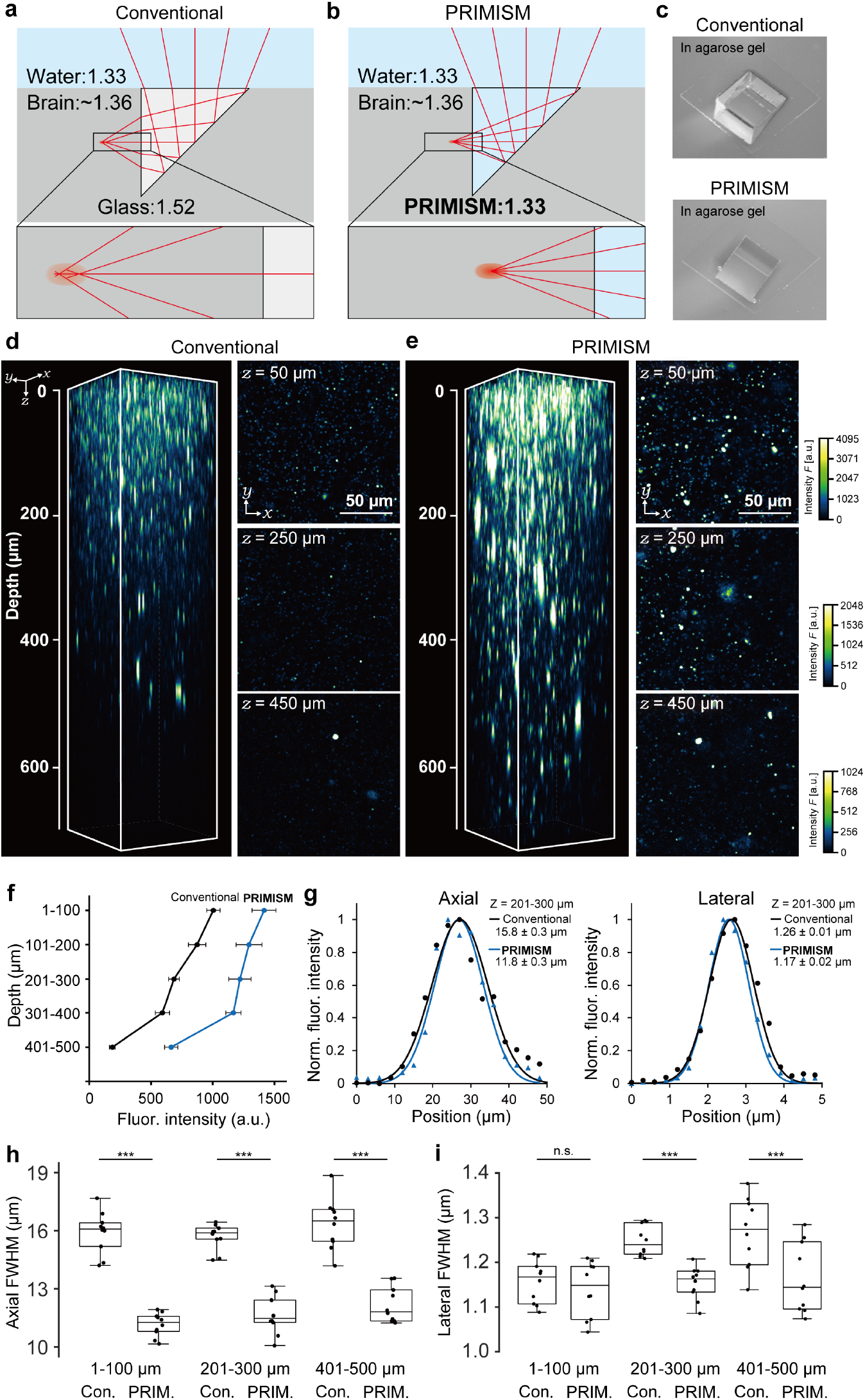
Optical performance evaluation using brain-mimicking gel model. (a, b) Ray-tracing schematics showing light paths and spherical aberration generated in a conventional glass prism (a) and the refractive index-matched PRIMISM (b). (c) Photographs of conventional glass prism and PRIMISM embedded in agarose gel. (d, e) Fluorescence images of 0.2-μm-diameter beads embedded in brain-mimicking gel, acquired through a conventional glass prism (d) or the PRIMISM (e). The left panel shows a 3D (xyz) reconstruction, while the right panel presents representative xy images at depths of 50 µm, 250 µm, and 450 µm. (f) Comparison of fluorescence intensity as a function of imaging depth for each prism type. (g) Representative point spread function (PSF) profiles at a 250-µm depth showing axial and lateral distributions. (h) Summary of axial PSF evaluations at 100-µm, 250-µm, and 400-µm depths for each prism type (n=10 beads/depth). FWHMs of PRIMISM: 11.2 ± 0.19 µm at 100 µm, 11.8 ± 0.26 µm at 250 µm, and 12.2 ± 0.29 µm at 400 µm; conventional glass prism: 15.9 ± 0.34 µm at 100 µm, 15.8 ± 0.28 µm at 250 µm, and 16.3 ± 0.42 µm at 400 µm; mean ± standard error of the mean. (i) Summary of lateral PSF evaluations (n=10 beads/depth). FWHMs of PRIMISM: 1.14 ± 0.02 µm at 100 µm, 1.17 ± 0.02 µm at 250 µm, and 1.16 ± 0.02 µm at 400 µm; conventional glass prism: 1.16 ± 0.02 µm at 100 µm, 1.26 ± 0.01 µm at 250 µm,, and 1.27 ± 0.02 µm at 400 µm; mean ± standard error of the mean.

Although the performance of the PRIMISM would ideally be assessed *in vivo*, such validation in mouse brains is challenging due to considerable individual variability in tissue structure, particularly the vascular anatomy, which greatly affects the prism implantation success rate. Therefore, to ensure more stable and quantitative evaluation, we investigated the optical properties of the PRIMISM by conducting *in vitro* two-photon imaging of a gel model mimicking the optical properties of brain tissue (i.e., with matching refractive index and scattering properties), composed of agarose, intralipid^31^, and tiny fluorescent beads (Fig. 1c, Supplementary Fig 3,4). Using this model, we demonstrated that the PRIMISM improves the brightness of the fluorescent images at all depths, compared to a conventional glass prism (Fig. 1d–f). We also estimated the lateral and axial resolutions by measuring the full-width at half maximum (FWHM) values of the point spread function (PSF), which was defined from the fluorescence intensity profile of a single bead smaller than the system’s resolution (Fig. 1g–i). The axial resolution was markedly improved when using the PRIMISM: we demonstrated significantly smaller axial FWHM values consistently across all depths in the brain-mimicking gel from the surface of the PRIMISM (Fig. 1g, i). These results indicate that the improved fluorescence intensity and spatial resolution are likely due to the improved photon collection efficiency achieved through reduced aberrations. Furthermore, Fig. 1d-f show that the PRIMISM may allow detection of signals that are undetectable by a conventional glass prism in deep regions (e.g., layer 5/6 during prism-based mPFC imaging). We also observed that the lateral resolution achieved with the PRIMISM was subcellular (substantially smaller than the neural cell body size, approximately 10 µm) and significantly improved, especially in deep regions (Fig. 1g, h). In addition, we evaluated chromatic aberration using tiny fluorescent beads and confirmed that no significant aberration occurred (Supplementary Fig. 4). This comparative analysis using brain-mimicking gel with conventional glass prisms demonstrated that the PRIMISM greatly improves *in vivo* optical imaging, making it a powerful tool for observing neural structures and functions with less distortion.

## *In vivo* high-resolution structural imaging in the deep mPFC

We next evaluated the performance of the PRIMISM in a living mouse brain by implanting it into transgenic mice expressing yellow fluorescent protein (YFP) in deep-layer excitatory cortical neurons^38^ and performing two-photon imaging of neural morphology in the mPFC (Fig. 2a–d). The PRIMISM-enabled high-resolution imaging of the mPFC neurons, even at depths exceeding 1.3 mm from the dorsal surface of the brain, clearly resolved subcellular structures such as dendritic spines (Fig. 2e, f). The PRIMISM allowed clear observation from the superficial to deep cortical layers in the deep mPFC (Fig. 2d–f), allowing for detailed *in vivo* observation of the deep neuronal architecture that was previously difficult to visualize.

**Figure 2.**
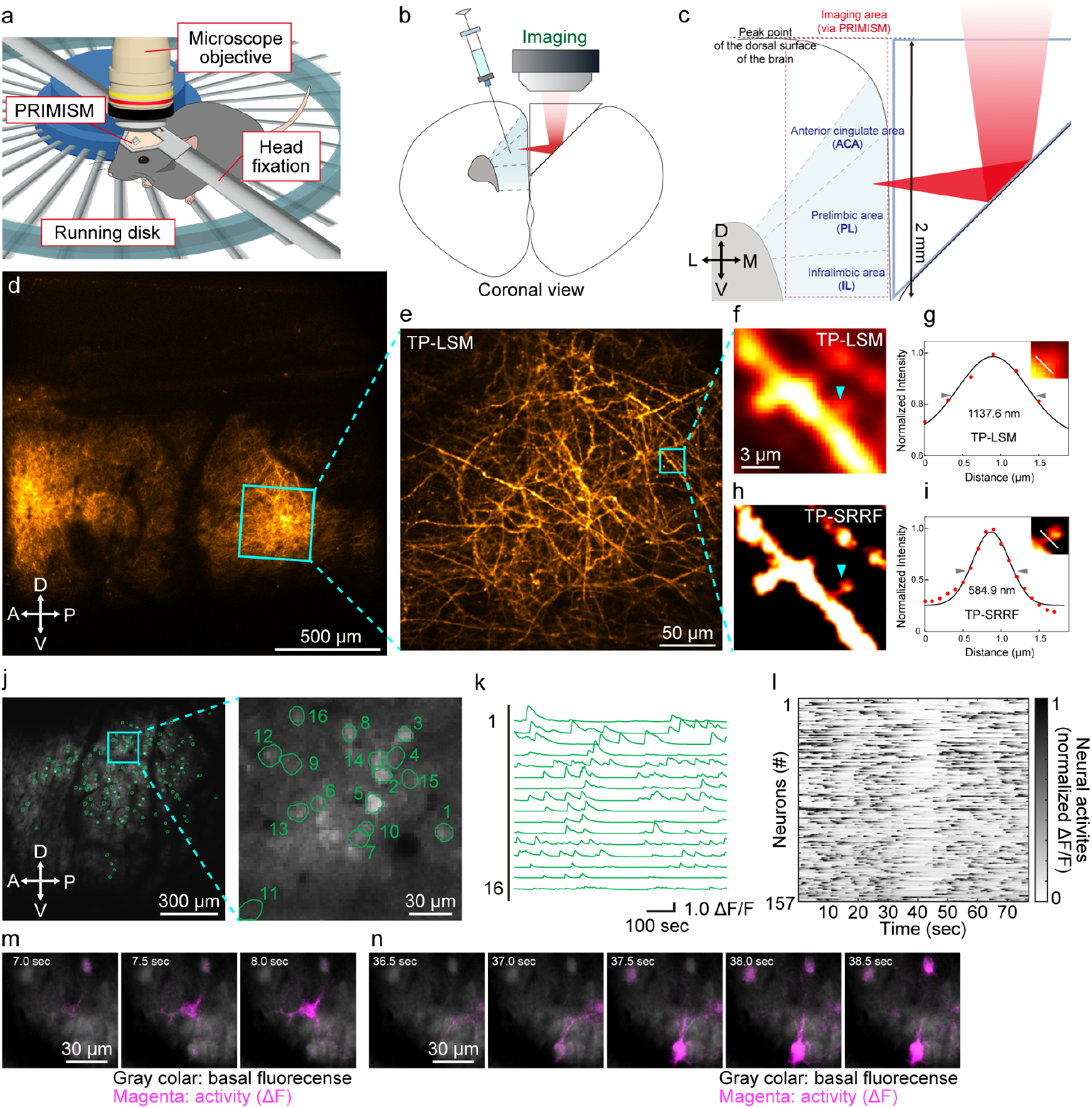
*In vivo* high-resolution deep-brain imaging using PRIMISM. (a) Schematic of the experimental setup for *in vivo* two-photon imaging with the PRIMISM in head-fixed mice. Structural imaging was performed under anesthesia, whereas neural activity imaging was conducted in awake mice. (b) Coronal view of the cranial window and PRIMISM insertion site along the midline fissure. PRIMISM implantation along the midline for optical access to the mPFC without transecting nerves. For the neural activity imaging, GCaMP6f was expressed in the mPFC excitatory neurons by AAV under CaMKII promoter regulation. (c) Schematic coronal section of the mouse brain showing the imaging plane and accessible mPFC areas: anterior cingulate area (ACA), prelimbic area (PL), and infralimbic area (IL). (d-i) Example of *in vivo* dmPFC imaging of Thy1-EYFP-H (H-line) transgenic mice implanted with the PRIMISM. (d)*In vivo* two-photon laser scanning microscopy (TP-LSM) image of the mPFC in a Thy1-YFP-H mouse acquired with the PRIMISM. (e) Higher magnification view of the mPFC region shown in (d). (f) Magnified view of the dendrite demarcated by the cyan box in (e). (g) Intensity profile of TP-LSM image along the white dashed line shown in the insert (taken from cyan arrowhead in (f)). (h) SRRF-processed image corresponding to the same dendrite depicted in (f). (i) Intensity profile of the TP-SRRF image along the white dashed line shown in the insert (taken from cyan arrowhead in (h)). (j-n) Example of *in vivo* mPFC imaging of GCaMP6f injected mice implanted with the PRIMISM. (j) Left: *in vivo* Ca^2+^ imaging of GCaMP6f-expressing neurons in the mPFC. Right: magnified view of the area demarcated by the cyan box in the left panel. Circles indicate ROIs. (k) Neural activity traces (ΔF/F) from 16 individual neurons, shown in the left panel of (j) (l) Raster plot of neural activity across all neurons shown in the right panel of (j). (m, n) Example time-lapse images of Ca^2+^ transients (magenta, ΔF) overlaid on baseline fluorescence (gray), demonstrating subcellular changes. D, dorsal; V, ventral; A, anterior; P, posterior; L, lateral; M, medial. Arrowheads next to the intensity profiles in (g) and (i) indicate the positions where the FWHM were calculated.

To further enhance spatial resolution, we applied the Super-Resolution Radial Fluctuations (SRRF) technique^39, 40^, an analysis-based resolution enhancement technique, to images of neural structures acquired through the PRIMISM. Using brain-mimicking gel, we first confirmed that the FWHM of fluorescent beads decreased from 1139.4 ± 21.7 nm to 502.3 ± 8.0 nm with SRRF near the surface of the PRIMISM, and from 1199.6 ± 13.1 nm to 556.3 ± 15.2 nm at a 300-µm depth (Supplementary Fig. 5a–c). A similar trend was observed under the conventional glass prism, but PRIMISM consistently yielded better resolution at both depths (Supplementary Fig. 5). Next, the effectiveness of the SRRF was also validated in neuronal tissue imaging: spine necks that appeared blurred in Fig. 2f were clearly resolved after SRRF processing (Fig. 2h), with FWHM values reduced from 1137.6 nm to 584.9 nm (Fig. 2g, i, and Supplementary Fig. 5d, e). Importantly, the SRRF analysis requires multiple rounds of time-lapse imaging with sufficient image quality (i.e., S/N ratio) to promote accurate reconstruction. Because the PRIMISM improves spatial resolution and brightness (Fig. 1), which in turn improves temporal resolution of the imaging as discussed above, the PRIMISM is expected to be advantageous in capturing the high temporal dynamics of such fine neural structures. Together, these results indicate that the PRIMISM combined with SRRF reliably achieves submicron resolution in the mPFC *in vivo* at depths exceeding 1.3 mm, facilitating precise visualization of the fine spine structures.

## Neural activity imaging across nearly the entire mPFC

To further demonstrate the PRIMISM’s utility for functional imaging, we captured real-time Ca^2+^ dynamics in neurons and dendrites within the mPFC of awake mice (Fig. 2a). Prior to the *in vivo* imaging experiments, mice were injected with an adeno-associated virus (AAV) expressing a genetically encoded Ca^2+^ indicator (GCaMP6f) throughout the mPFC, and the PRIMISM was subsequently implanted beside the injected area (Fig. 2b, c). Consistent with the high spatial-resolution demonstrated in the brain-mimicking gel experiments, *in vivo* changes in Ca^2+^ signals were clearly detected from fine neuronal structures, including cell bodies (Fig. 2j–l) and dendrites (Fig. 2m, n). We successfully extracted neural activity signals (i.e., Ca^2+^ transients) at cellular resolution across a wide FoV (Fig. 2j and Supplementary Video 1). The temporal dynamics of Ca^2+^ signals from representative neurons, where independent neural activity traces were clearly extracted, are shown in Fig. 2k, highlighting the ability to obtain single-cell resolution across a wide FoV. The raster plot shown in Fig. 2l demonstrates the ability to identify a large population in the FoV. Even when we expanded the FoV to cover the full FoV through the PRIMISM window, the enhanced brightness and resolution provided by the PRIMISM allowed for sufficient S/N ratios, even at the lateral edges where detection is typically compromised, providing for comprehensive neural activity monitoring across the entire mPFC at single-cell resolution (Supplementary Fig. 7 and Supplementary Video 2). The PRIMISM also facilitated the detection of localized Ca^2+^ transients within fine neuronal structures, such as dendrites (Fig. 2m, n and Supplementary Video 3). These time-lapse sequences illustrated dynamic fluorescence changes (magenta) over a basal background (gray), confirming subcellular activity detection with high spatiotemporal resolution. Furthermore, as demonstrated in the brain-mimicking gel experiments, PRIMISM is advantageous for imaging along the depth axis. Thus, we demonstrated *in vivo* four-dimensional (4D) Ca^2+^ imaging, capturing volumetric activity dynamics across multiple depths within the mPFC at cellular resolution (Supplementary Fig. 8 and Supplementary Video 4).

These capabilities are particularly valuable for investigating the spatiotemporal dynamics of neuronal signaling and network interactions. The PRIMISM thus provides a powerful platform for chronic functional imaging at both fine and wide spatial scales.

## Discussion

The PRIMISM allows for substantial progress in deep-brain imaging, enabling bright, high-spatial-resolution visualization of neuronal structures and activity at depths exceeding 1.3 mm from the brain surface.

The reduced optical aberrations enhance the PRIMISM’s compatibility with advanced light-shaping strategies, providing another compelling pathway for expanding its utility. Such strategies include adaptive optics using spatial light modulators to further improve spatial resolution. These technologies show great potential in correcting aberrations arising from refractive index mismatches and tissue heterogeneity in *in vivo* two-photon microscopy^35, 41-43^. Combining these technologies with the PRIMISM could enable deeper, brighter, and more spatially uniform imaging across wide FoVs and unlock new applications in precise functional interrogation of neural circuits. Additional approaches include Bessel beam scanning for fast volumetric imaging and holographic photostimulation for multi-point optogenetic manipulation.

Furthermore, the PRIMISM’s compact and modular design makes it a promising candidate for future integration into miniscope imaging systems for freely moving animals^24^, offering the potential for long-term imaging of deep-brain activity during natural behavior. While a previous study combined a conventional glass prism with two-photon microscopy for deep-brain imaging in freely moving mice^25^, further improvements in optical performance may be achievable with our PRIMISM technology. Additionally, combining the PRIMISM with GRIN lens systems may provide new hybrid approaches for deep imaging, especially in miniaturized settings. The potential to combine the PRIMISM with lightweight optical components, calcium indicators, and wireless recording technologies could open new avenues for studying dynamic brain functions under ethologically relevant conditions.

Future directions for the PRIMISM include its application to *in vivo* multicolor, time-lapse super-resolution imaging, enabling the visualization of subcellular dynamics at high temporal resolution. Recent advances in four-dimensional imaging of live multicellular organisms^44-46^ have highlighted the potential of such techniques to uncover fundamental biological processes in the vertebrate brain as well.

In summary, the PRIMISM represents a substantial advancement in deep-brain optical imaging, providing a refractive index-matched interface that enhances both spatial resolution and brightness. These improvements enable wide-field two-photon imaging across multiple cortical layers *in vivo*, facilitating comprehensive visualization of neural structures and dynamics at subcellular resolution. Its successful application to high-resolution imaging of the mPFC— including the SRRF and *in vivo* Ca^2+^ dynamics—demonstrates its versatility for probing neural circuits, cognitive functions, and disease-related processes. With further refinement and integration with emerging optical technologies, PRIMISM has the potential to become a foundational platform for next-generation neuroscience and biomedical research.

## Methods

### Animals

All animal experiments were carried out in accordance with the Institutional Guidance on Animal Experimentation and with permission from the National Institutes of Health guidelines and approved by the National Institute for Physiological Sciences Animal Care and Use Committee. All mice were housed under a 12 h/12 h light/dark cycle with free access to food and water in a temperature-controlled environment (22–24°C and 30–60% humidity). Adult wild-type C57BL/6 J mice (males and females over 8 weeks of age) were used for intracellular calcium imaging. Thy1-EYFP-H (H-line)^38^ transgenic mice were used for imaging of neuronal morphology.

### Fabrication of PRIMISM

The mirror was fabricated by coating a cover glass with aluminum. A 0.06-0.17 mm-thick (Matsunami cover glass No. 00) cover glass was cleaned by applying an optical cleaner (Photonic Cleaning Technologies, First Contact). Prior to the aluminum deposition, the surface of the cover glass was activated with oxygen plasma cleaner (Pie Scientific, Tergeo Plus plasma cleaner) at 5 standard cubic centimeters per minute (sccm) O_2_ and 150 W for 5 min to improve the adhesion of the aluminum to the cover glass. A 300-460 nm-thick aluminum coating was then thermally evaporated at 50 Å/s on the cover glass. The mirror was trimmed to 1.9 mm x 2.5 mm using a UV laser (Keyence, MD-U1000C) while protecting the aluminum layer from dust with optical cleaner. The mirror, low-refractive index UV-curable polymer (MY Polymers, BIO-133), and another cover glass (Matsunami cover glass No. 00) were placed on a metal mold (Supplementary Figs. 1, 2). For ease of handling, the mold comprised five parts. Gaps between parts of the mold were minimized by tightening the screws from each of the sides. The sample in the mold was exposed to 275 nm UV by UV-LED for 11 min to cure the polymer. The width of the mold for the PRIMISM was designed to be larger than the width of the mirror to prevent bubble formation in the optical path used for imaging. A bubble was inevitably formed due to the polymer shrinking during the curing, but the wider mold design allowed the bubble to form in the peripheral region, outside of the optical path. Prior to the photo-curing, the surface of the mirror and cover glass were activated using oxygen plasma cleaner (Pie Scientific, Tergeo Plus plasma cleaner) at 5 sccm O_2_ and 50 W for 1 min to improve adhesion with the polymer. Fabrication of the PRIMISM was completed after removing the sample from the mold.

### Fluorescent bead imaging and analyses

To evaluate the spatial resolution of PRIMISM imaging, yellow-green fluorescent 0.2-µm diameter beads (FluoSpheres, Carboxylate-modified, [505/515], Invitrogen) embedded in 2% w/v agarose gel (agarose L, Nippon Gene) or the brain-mimicking gel with optical properties (scattering coefficient =10 cm^−1^ and refractive index=1.36) similar to those of living mouse brain cortex^39^ were used. Before the gel solidified, a PRIMISM or a glass prism was placed on the gel. To observe the fluorescent beads, a water-immersion ×10/0.5 NA objective lens (CFI Plan Apo 10XC Glyc, Nikon) was used with a two-photon excitation microscope (A1R-MP+, Nikon). Image stacks were acquired with *xy* pixel size of 300 nm and z-steps of 3 µm at 960 nm of Ti-Sa excitation laser (MaiTai eHP DeepSee, Spectra Physics). The xy-and z-FWHM values of the PSFs were calculated by Gaussian curve-fitting of the fluorescence intensity profiles of each bead image using ImageJ.

To evaluate the axial chromatic aberration of the PRIMISM, the same samples used for spatial resolution analyses were observed while switching the excitation wavelength of the Ti-Sa laser to 800, 900, and 1000 nm. Fluorescent images were then acquired via two separate color channels, 500-550 nm and 562.5-587.5 nm. The line profiles along the *xz*-axis of each single fluorescent bead were measured using NIS-Elements AR analysis software (ver. 5.20.00, Nikon) and exported in CSV format. The line profiles were then fitted by a Gaussian function, and the differences in the center positions of the peaks between different color channels were calculated.

### Virus injection

To express GCaMP6f, a genetically encoded calcium indicator for monitoring neural activity, we used a gene expression system based on the AAV vector, as described previously^4, 5^. Viruses were injected into mice at postnatal day (P) 50–190 for *in vivo* experiments, at least 3 weeks before microprism implantation, which was followed by *in vivo* experiments at least 1 month after the implantation. During the surgery for the virus injection, the mice were anesthetized with isoflurane (induction 2% [partial pressure in air] and then reduced to 1%). A small square (∼1 × 1 mm^2^) of the skull was removed over the left mPFC using a dental drill to mark the site for a small craniotomy. AAV1/CamKII.GCaMP6f was obtained from the Addgene (#100834-AAV1), and injected into the left or right mPFC at three sites (depth 1.0, 1.5, and 2.0 mm from the dorsal surface of the brain, volume 375 nl/site of 1:1 diluted aliquot (originally 2×10^13^ GC/ml) at 75 nl/min) to cover the mPFC area, over a 5-min period at each depth using a UMP3 microsyringe pump (World Precision Instruments), NanoFil 10-µl Syringe (World Precision Instruments) and Nanofil 35G Beveled Needle (World Precision Instruments). Because the large blood vessels stemming from the sinus interfere with the insertion of the PRIMISM, the AAV injection was performed in the hemisphere contralateral to the region with a wide-open area (i.e., free of large blood vessels). The X-Y coordinates of the injection site were usually 0.5 mm lateral to the midline and 2.0 mm rostral from bregma, but if large blood vessels obstructed the position, the insertion site was slightly shifted to avoid the vessels. The beveled side of the injection needle faced the midline so that the needle could be smoothly inserted and the virus would cover the mPFC. We carefully designed our injection protocol (especially the volume and depth) to cover almost the entire mPFC, which includes, according to the nomenclature in the Allen Brain Atlas (https://atlas.brain-map.org/), the ACA (anterior cingulate area), PL (prelimbic area), and the IL (infralimbic area) (Fig. 2c).

### *In vivo* two-photon imaging

*In vivo* two-photon imaging was performed as described previously^4-6,30^, with modifications to pair with our new experiments based on the PRIMISM.

For imaging GCaMP6f signals *in vivo* at 3 weeks after the virus injection, the mice were anesthetized with isoflurane (induction 2% [partial pressure in air] and reduced to 1%). A custom-made stainless steel (or aluminum) head plate was attached to the skull with dental cement (Super-Bond, Sun Medical) as shown in Fig. 2a and Supplementary Fig. 6a. For the subsequent PRIMISM implantation, a square cranial window (approximately 2.8 × 2.8 mm^2^) was carefully made with minimal bleeding above the mPFC in the hemisphere opposite to the virus injection site. An implantable microprism assembly (Supplementary Fig. 2), comprising the PRIMISM bonded to a 3 × 4 mm cover glass (No. 00; Matsunami), was prepared and inserted into the subdural space within the fissure along the midline, as described previously^4, 30^ to avoid harming any nerves surrounding the mPFC network in both hemispheres, thereby allowing for visualization of the AAV-injected side of the mPFC through the imaging window. The area directly beneath the microprism was compressed but remained intact. This insertion procedure sometimes caused a small amount of bleeding that covered the imaging site, but even in those cases, the imaging window became clear after waiting at least 3 weeks before performing the imaging experiments. As reported before^4, 30^, the mice recovered quickly and displayed no gross impairments or behavioral differences compared with non-implanted mice, enabling chronic imaging of the mPFC in behaving mice.

The anatomic coordinates of the FoV for two-photon imaging within the mPFC were precisely measured using the position of the dorsal surface of the brain, the sinus, and the pial surface along the midline, which were usually visible through the imaging window prepared as described above, as a guide. The depth from the dorsal surface of the brain and the distance from the surface of the mPFC (i.e., the end of the PRIMISM) of the imaging area within the mPFC were recorded as described previously^47^. In addition to the mPFC, neural activities were recorded through the implanted PRIMISM in the ACA, PL, and IL, from the superficial layer (L1) to the deep layer (L5/6) (∼600 μm from the pial surface along the midline) of the dmPFC (Fig. 2c).

Activities of mPFC neurons were recorded by imaging GCaMP6f fluorescence changes with a FVMPE-RS two-photon microscope (Olympus) and a Mai Tai DeepSee Ti:sapphire laser (Spectra-Physics) at 920 nm, through a 4× dry objective, 0.28 N.A. (XLFLUOR4X, Olympus) or a 10× water immersion objective, 0.50 N.A. (CFI Plan Apo 10XC Glyc, Nikon), as described previously^4^. The microscope is capable of 30-Hz scans (per z plane) at 512 × 512-pixel resolution and even higher temporal resolution with smaller pixel sizes. It will be crucial to optimize the scanning parameters by switching between “four-dimensional (4D) volumetric scanning” for broad area screening and “small-volume or single z-plane imaging” for detailed measurements with faster temporal resolution. The temporal resolution for each experiment in the present study is provided in the corresponding figure legend. GCaMP6f signals were detected via the band-pass emission filter (495–540 nm). As the GCaMP6f was expressed under regulation of the CaMKII promoter^48, 49^, all of the recording targets were assumed to be excitatory neurons^50^. Scanning and image acquisition were controlled by FV30S-SW image acquisition and processing software (Olympus). To smoothly position the mice below the objective lens for imaging, light and minimal-duration isoflurane (2.0% for <2–3 min) anesthesia was used. Behavioral and imaging experiments were started 5 mins after the mice recovered from the anesthesia and began locomoting on a running disk. Locomotor activity was visually confirmed using a video camera (VLG-02, Baumer) under infrared light-emitting diode illumination (850 nm: LDL-130×15IR2-850, CCS Inc.).

To evaluate the performance in imaging neural and synaptic structures, fluorescence images of the Thy1-EYFP-H (H-line) mice (The Jackson Laboratory, Strain #:003782) were obtained using a two-photon laser microscopy customized for *in vivo* imaging (A1R-MP+, Nikon), as described previously^6^. The Nikon CFI Plan Apo 10XC Glyc (10x, 0.5 NA, water-immersion objective lens) was used for most of the experiments (e.g., Fig. 2d-i), whereas the Olympus XLFLUOR4X/340 (4x, 0.28 NA, air-immersion objective lens) was used to obtain images with an entire FoV through the PRIMISM (e.g., Fig. 2d). A Ti:Sapphire laser (MaiTai eHP DeepSee, Spectra Physics) was used as an excitation light source. EYFP images were acquired using an excitation wavelength of 950 nm and a 560-nm dichroic mirror with a highly sensitive GaAsP-NDD detector. Scanning and image acquisition was controlled by NIS-elements (Nikon).The EYFP images were obtained from mice under anesthesia (isoflurane, 1.0–1.5%). Body temperature was maintained during imaging using a disposable heating pad.

### Imaging data analyses (and statistics)

Raw images of the GcaMP6f signals in the mPFC were processed to correct for brain motion artifacts using the enhanced correlation coefficient (ECC) image alignment algorithm^51^, as described previously^5, 52^. The regions of interest (ROIs) for the detection of neural activity were automatically selected using a constrained nonnegative matrix factorization algorithm in MATLAB (versions: R2019b, R2022a, R2023b) as described previously^53^, with some manual adjustment. Further steps to process the GcaMP6f signals for measurements of the signal change (ΔF/F) from each neuron were performed as described previously^5, 52, 54^; the same constrained nonnegative matrix factorization package for ROI detection (i.e., extraction of active neurons) also provides an option for signal processing, which was employed in the present study to calculate ΔF/F for each selected neuron.

In the case of volumetric live imaging, time-lapse images at various depths were acquired by sequentially adjusting the imaging focal plane (Supplementary Fig. 8). Motion artifacts in the time-series of GCaMP images at each z-plane were corrected using the ECC image alignment algorithm. Following ROI extraction of active neurons at each z-plane, as described above, we assessed the spatial overlap of the ROIs across z-planes. For this purpose, motion-corrected images from each z-plane were averaged over time, and the obtained average images were further aligned across z-planes to correct for the shift between z-planes using the ECC image alignment algorithm^51^. For this correction along the z-axis, the first z-plane (i.e., the most superficial one) was used as a reference image for correcting the second z-plane. Then, an average image of these two aligned z-planes was further used for correction of the third z-plane. The average image of these three aligned z-planes was then used for the motion correction of the fourth z-plane. Through this procedure, four z-planes were motion corrected and used to accurately check the overlap of the extracted ROIs across z-planes.

### SRRF processing

The NanoJ-eSRRF plug-in (ver. 1.1.0) of Fiji/ImageJ (ver. 1.54h) was used for eSRRF, an image-analysis-based super-resolution process^55^. Processing was performed using 30 consecutively acquired images. The parameters of the eSRRF analysis were set as follows: Magnification = 5, Radius = 4.00, Sensitivity = 4. The AVG reconstruction mode was used for the image reconstructions.

## Author Contributions

M.A. conceived and coordinated the whole project with T.T, K.O., and N.M.; T.T., Y.Z., K.O., N.M., and M.A. designed and developed a method for PRIMISM with the support of M.T., J.N., and T.N.; T.T., M.T., and M.A. performed *in vitro* imaging experiments using the PRIMISM; T.T., C.I., and M.A. performed *in vivo* imaging experiments using PRIMISM with the support of A.H., H.I., A.S., H.Y., J.N. and T.N.; T.T., M.T., A.H., and M.A performed data analyses; T.T., K.O., N.M., and M.A. wrote the paper, with contributions from all authors.

## Data availability

Data supporting the findings of the present study are available from the corresponding authors upon reasonable request. Anatomic information in the Allen Brain Atlas (https://atlas.brain-map.org/) was used for the anatomic description, determination of the virus injection area, and evaluation of the recorded brain regions.

## Acknowledgements

We thank the members of the laboratory for their help, especially Yuki Watakabe, Ryoko Arakawa, and Aki Hiraoka for experiments. We also thank Arclev Academia Strategists Network for providing a collaborative environment that facilitated the initial interactions leading to this interdisciplinary research.

