## Supplemental figures for "*In vivo* deep-brain microscopy at submicrometer resolution with refractive index-matched prism interfaces"

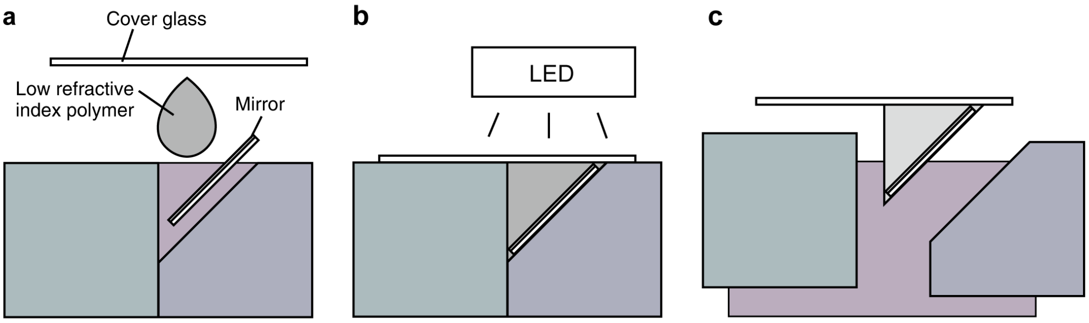


**Supplementary Fig. 1. PRIMISM fabrication.**

(a) Put a piece of mirror, polymer, and a piece of cover glass into the mold.

(b) Cure with 275 nm LED light.

(c) Remove PRIMISM from the mold.

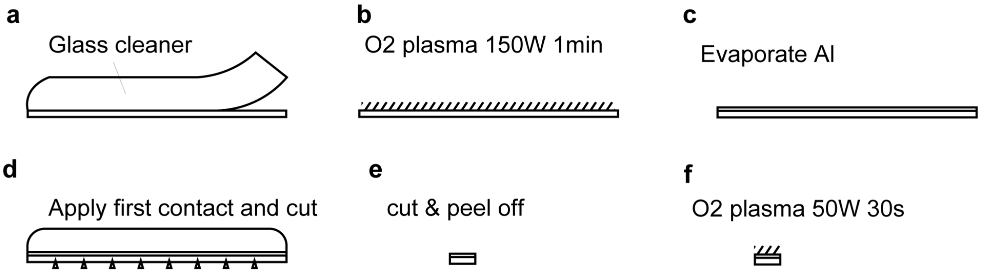


**Supplementary Fig. 1-1. Mirror fabrication.**

(a) First Contact was applied on and peeled off from the cover glass.

(b) Oxygen plasma was applied at 150 W for 5 min to increase the adhesion of the evaporated aluminum.

(c) Vacuum evaporation of the aluminum, at 50 Å/s for 340 - 400 nm.

(d) First contact was applied on the aluminum side, then the laser cut was applied from the glass side. The mirror size was 1.9 mm by 2.5 mm.

The cover glass was cut and peeled off from the First Contact film.

(f) Oxygen plasma was applied just before the fabrication of the PRIMISM.

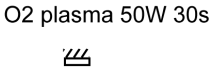


**Supplementary Fig. 1-2. Fabrication of the cover glass used in PRIMISM.**
A piece of cover glass (3 mm by 4 mm) was prepared. Then, oxygen plasma was applied at 50 W for 30 s to increase the adhesion of the polymer.

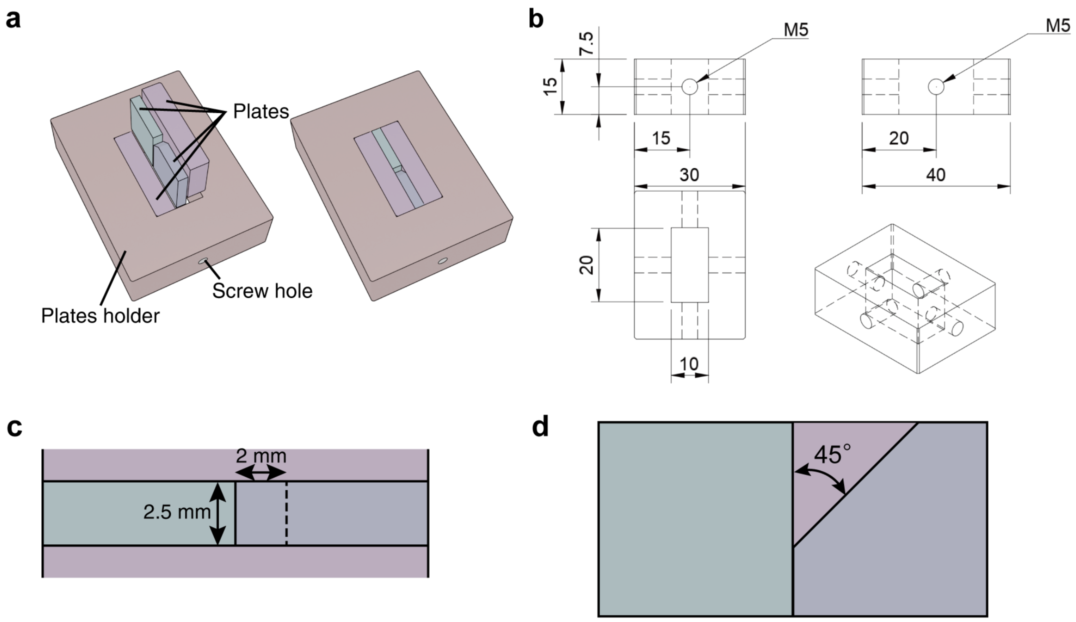


**Supplementary Fig. 2. The mold for the PRIMISM.**

(a) The PRIMISM mold is consisted of 4 plates and 1 plate holder. The plates are fixed with 4 screws to the plate holder.

(b) The measurements of the plate holder. (unit: mm)

(c) Top view of the fixed plates. X is the width of the PRIMISM, and Y is the size of the PRIMISM.

(d) Angle of the plates.

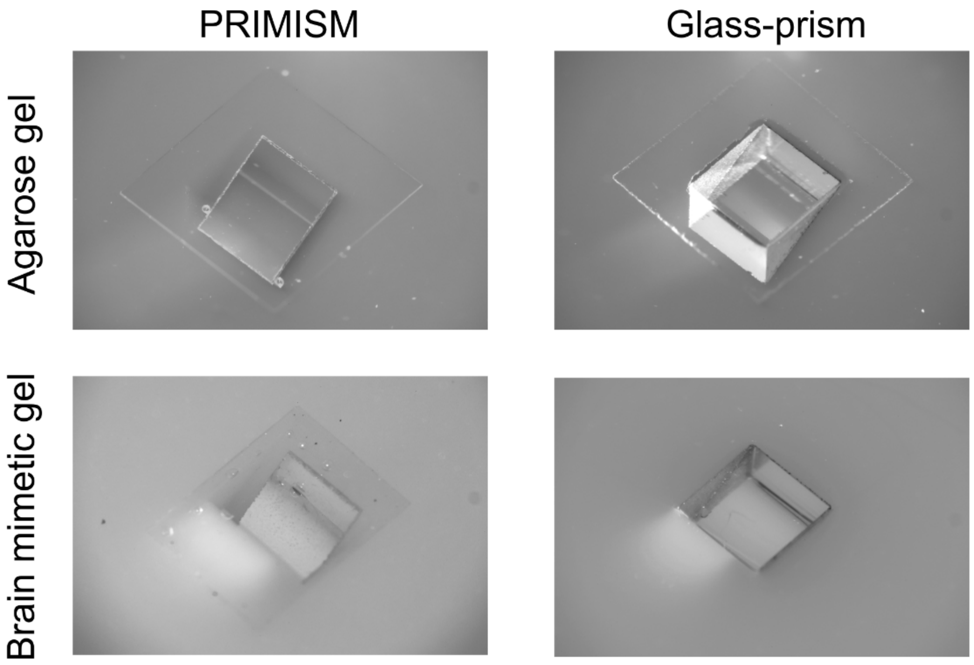


**Supplementary Fig. 3. PRIMISM and Glass-prism embedded in agarose gel or brain mimetic gel.**

Photographs taken from diagonally above of the prisms. The shooting angles and imaging conditions were matched as much as possible.

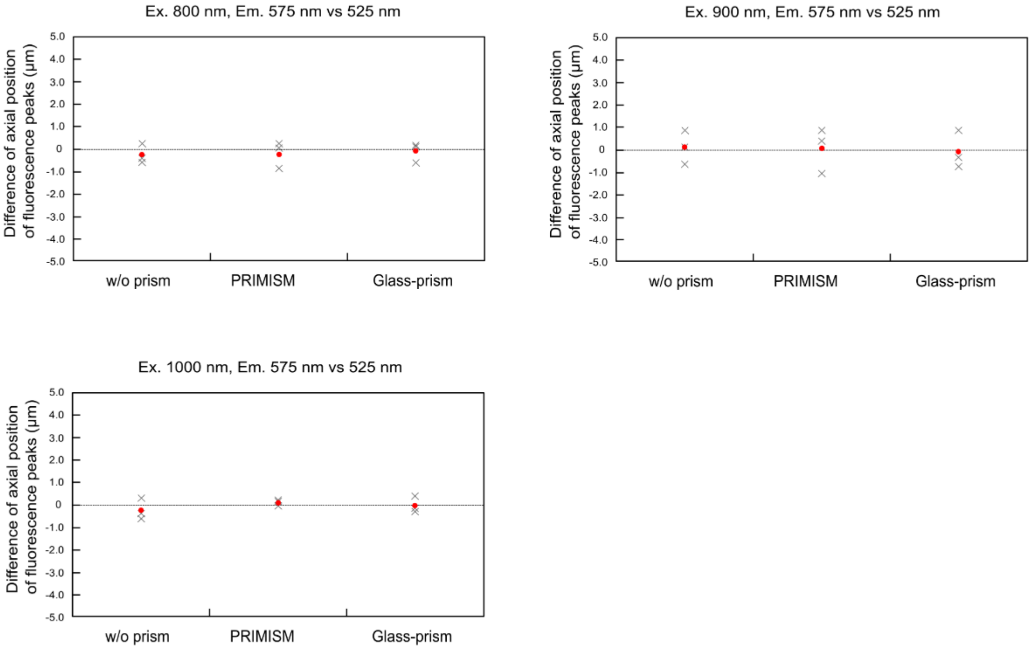


**Supplementary Fig. 4. Evaluation of axial chromatic aberration through the PRIMISM using fluorescent beads embedded in agarose gels at different excitation wavelengths.**

In each condition, a single fluorescent bead was observed by two-different fluorescence emission ranges; one near 525 nm (500-550 nm) and the other near 575 nm (562.5-587.5 nm). The peak positions of their fluorescence were determined by curve fitting of the intensity profiles using a Gaussian function, and the difference was calculated. The cross-marks show raw data plots ($n = 3$), and the red filled circles show the averages of the data.

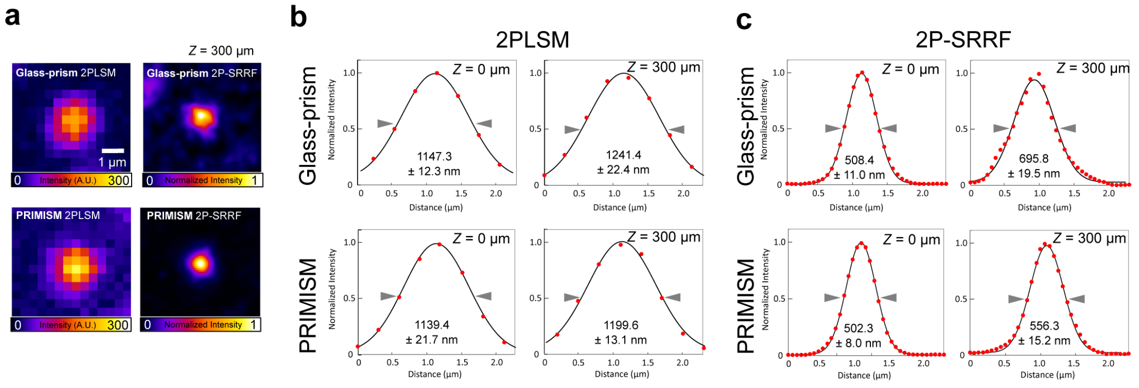


**Supplementary Fig. 5. Effect of PRIMISMS for super-resolution imaging using SRRF.**

(a) Fluorescent bead images observed by two-photon laser scanning microscopy (2PLSM) and two-photon super-resolution microscopy (2P-SRRF) at 300 μm depth in the brain mimetic gel under the glass-prism and PRIMISM. 2PLSM images show the average of consecutive images for SRRF process; 2P-SRRF images show the result of SRRF process from the same images.

(b) Representative intensity profiles of the fluorescent bead images of 2PLSM observation under glass-prism and PRIMISM at prism surface and 300 μm depth. The profiles show the intensity distribution in the *X*-direction at the center of the fluorescent beads. Black curves show the curve fitting using Gaussian function.

(c) Representative intensity profiles of the fluorescent bead images of 2P-SRRF observation under glass-prism and PRIMISM at prism surface and 300 μm depth. The values under each profile show the average and SEM of FWHM calculated from the analyses of 10 fluorescent beads in individual conditions.

Arrowheads next to the intensity profiles in (b) and (c) indicate the positions where the FWHM were calculated. The values under each profile show the average and SEM of FWHM calculated from the analyses of 5 fluorescent beads in individual conditions.

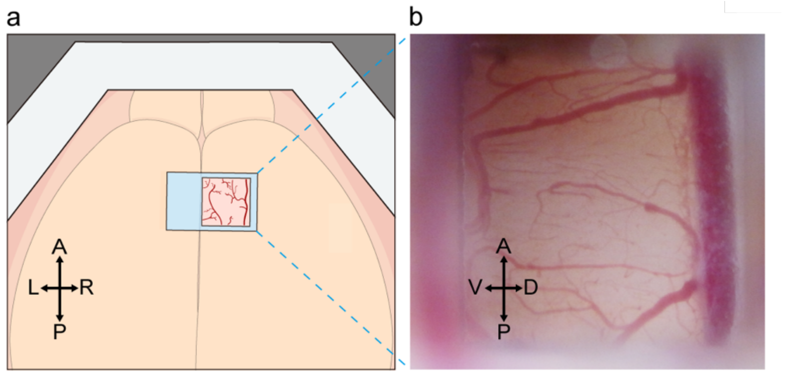


**Supplementary Fig. 6. PRIMISM implantation for chronic *in vivo* imaging of the mouse mPFC.**

(a) Schematic illustration showing the dorsal view of mouse brain, within the head plate fixed on the skull, after the PRIMISM implantation.

(b) Representative bright-field image of the mPFC observed through the PRIMISM window, one month after implantation, demonstrating the stable transparency across the entire field of view. Data from the same mouse are shown in Fig. 2j-n, Supplementary Figs. 7 and 8, and Supplementary Videos 1-4.

D, dorsal; V, ventral; A, anterior; P, posterior; L, left; R, right.

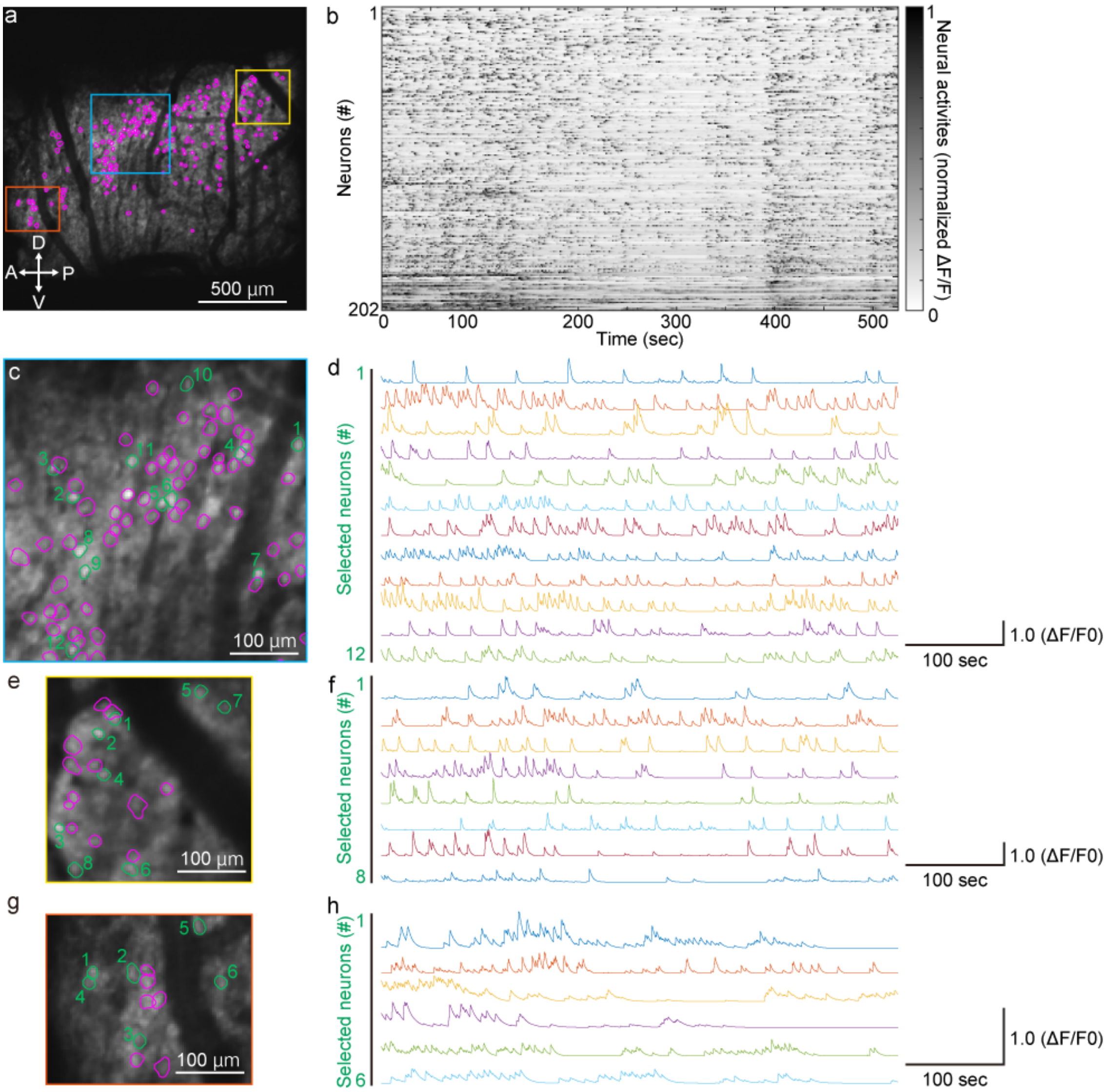


**Supplementary Fig. 7. PRIMISM enables wide-field two-photo imaging of mPFC neural population activity at cellular resolution.**

(a) Two-photon image showing GCaMP6f fluorescence in the mPFC of a mouse implanted with PRIMISM. Each region of interest (ROI) outlined in magenta corresponds to a putative single neuron. The baseline fluorescence image (gray) was calculated as median fluorescence across the recording session. These images were obtained from the same mouse as in Figure 2j and were acquired using a lower magnification objective.

(b) Normalized GCaMP6f ΔF/F signals over time for all ROIs shown in (a).

(c-h) Magnified views of the areas indicated by colored squares in (a): blue (c), yellow (e), and red (g). In each region, selected ROIs outlined in green correspond to neurons whose GCaMP6f traces are shown in (d), (f), (h), respectively.

See also Supplementary Video 2.

D, dorsal; V, ventral; A, anterior; P, posterior.

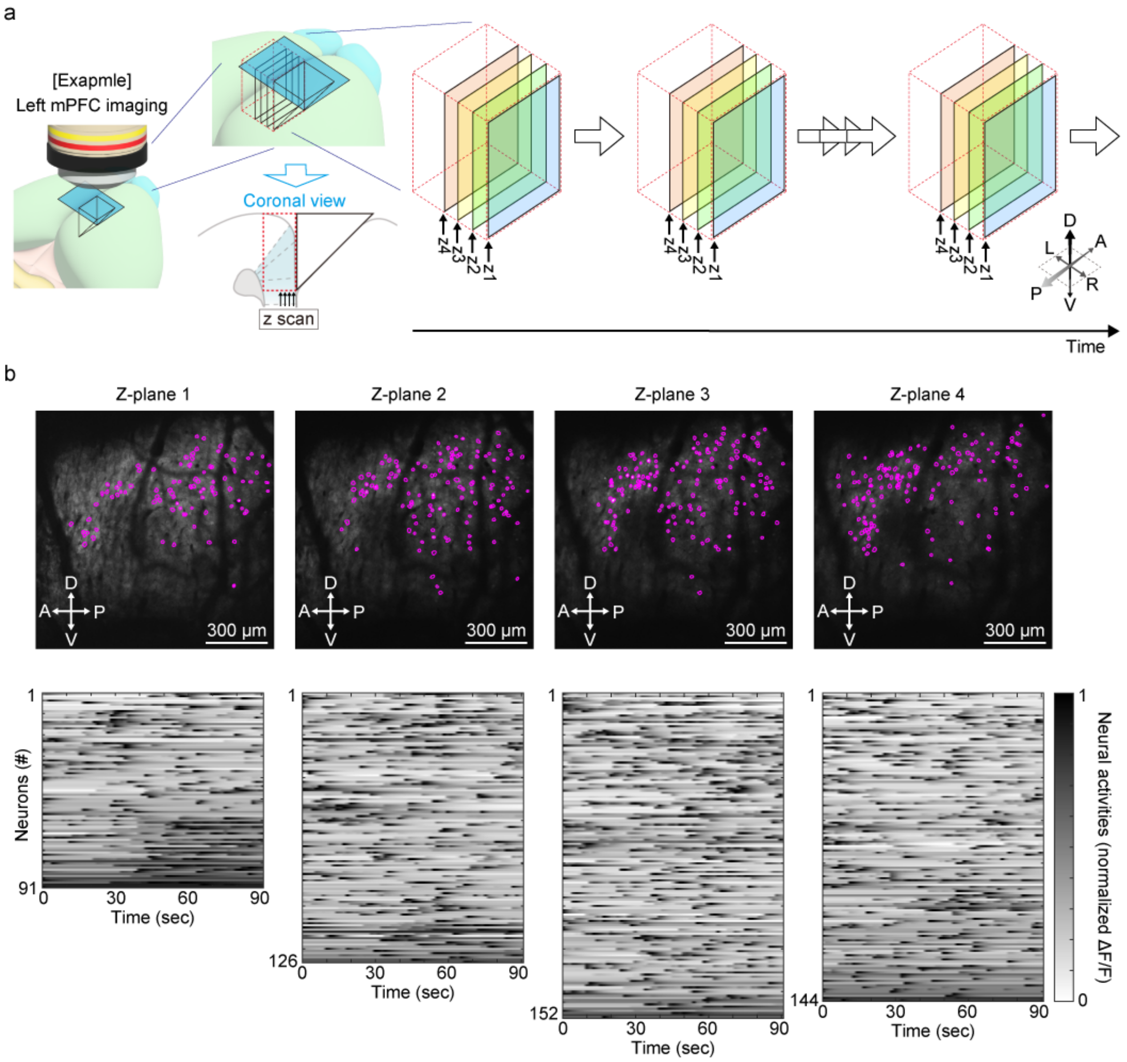


**Supplementary Fig. 8. Volumetric time-lapse imaging of mPFC neural activity**

(a) Schematic of *in vivo* two-photon time-lapse imaging at multiple depths through the PRIMISM implanted along the midline adjacent to the mPFC. Colored planes indicate the scanned z-planes (z1–z4).

(b) Representative *in vivo* volumetric imaging of the mPFC through the PRIMISM, from the same mouse shown in Figure 2j and Supplementary Figure 7. Four z-planes were acquired at each time point. (Top) Baseline fluorescence (gray) was estimated as the median fluorescence over time for each z-plane. ROIs corresponding to putative neurons are outlined in magenta. (Bottom) Grayscale heatmaps showing neuronal responses (normalized GCaMP6f signal) for each z-plane. See Methods for details. See also Supplementary Video 4.

D, dorsal; V, ventral; A, anterior; P, posterior; L, left; R, right.
